# SeqPanther: Sequence manipulation and mutation statistics toolset

**DOI:** 10.1101/2023.01.26.525629

**Authors:** James Emmanuel San, Stephanie Van Wyk, Houriiyah Tegally, Simeon Eche, Eduan Wilkinson, Aquillah M. Kanzi, Tulio de Oliveira, Anmol M. Kiran

## Abstract

Pathogen genomes harbor critical information necessary to support genomic investigations that inform public health interventions such as treatment, control, and eradication. To extract this information, their sequences are analysed to identify structural variations such as single nucleotide polymorphisms (SNPs) and insertions and deletions (indels) that may be associated with phenotypes of interest. Typically, this involves generating a consensus sequence from raw reads, aligning it to a reference and identifying positions where variations occur. Several pipelines exist to map raw reads and assemble whole genomes for downstream analysis. However, there is no easy to use, freely available bioinformatics quality control (QC) tool to explore mappings for both positional codons and nucleotide distributions in mapped short reads of microbial genomes. To address this problem, we have developed a fast and accurate tool to summarise read counts associated with codons, nucleotides, and indels in mapped next-generation sequencing (NGS) short reads. The tool, developed in Python, also provides a visualization of the genome sequencing depth and coverage. Furthermore, the tool can be run in single or batch mode, where several genomes need to be analysed. Our tool produces a text-based report that enables quick review or can be imported into any analytical tool for upstream analysis. Additionally, the tool also provides functionality to modify the consensus sequences by adding, masking, or restoring to wild type mutations specified by the user.

**Availability:** SeqPanther is available at https://github.com/codemeleon/seqPanther, along with the necessary documentation for installation and usage.

## Introduction

Next Generation Sequencing (NGS) platforms underlie an exciting era that facilitates the large-scale investigation of pathogen genomics. This in turn supports important aspects relating to public health including genomic epidemiological, pathogen surveillance, and pharmaceutical development of infectious diseases^1^. Despite these advances, quality control (QC) of sequence remains a concerning bottleneck to process big data in a timely fashion to generate actionable information. High quality and accurate genome assembly information provides insight on the occurrence and associated consequences of genetic changes within pathogenic microorganisms. Analyses to extract this information primarily involve the identification of variations such as single nucleotide polymorphisms (SNPs), and insertions and deletions (indels) which may be associated with enhanced pathogen phenotypes, such as increased transmissibility, immune and vaccine escape, increased infectivity, and disease severity^1^. For SARS-CoV-2, and other pathogens of public health interest, several pipelines exist to map sequenced raw reads to a reference genome and generate the consensus sequence for downstream analyses ^2–4^. However, often the consensus deviates from what is expected, requiring additional quality control (QC) and refinements prior to publication.

Additionally, in order to track microbial and viral diversity, genomic sequences are classified into lineages comprising a constellation of mutations exclusive to the lineage^5^. The absence of one or more lineage defining mutations, some of which may have been implicated in the altering of viral phenotypes, calls for further investigation. This includes re-examining the sequenced and mapped reads to establish the underlying reasons for its absence, and occasionally, and if warranted restore it.

Wrong and / or uncalled mutations, representing false positives and negatives, could arise due to several factors negatively affecting the sequencing, mapping, and assembly outcomes. These include primer drop-outs and algorithmic issues^6^. Algorithmic issues occur when expected parameter values significantly differ from the values encountered, for example, when depth of coverage is lower than the expected minimum depth due to low viral loads^7^. Low viral loads are typical of samples collected at the later stages of infection or tail ends of an outbreak commonly characterised by high cycle thresholds (Ct) values (> 30) or low viral loads^8^. Sequences from these are usually characterised by large numbers of frameshifts, indels, clustered and private mutations, i.e. mutations that are unique to a strain compared to their nearest neighbour in the global phylogeny for supported pathogens^9^. Primer drop-outs, on the other hand, are often caused by hypermutation in primer binding regions, reducing the amplification and sequencing for the targeted regions. This results in sections of the genome with low or no coverage. Primer drop-outs were commonly reported multiple times throughout the SARS-CoV-2 pandemic, especially in the variants of concern (VOCs)^8,9,10^. Mutations that have resulted in primer drop-outs for VOCs include the G142D (Delta and Omicron) in the 2_Right primer, the 241/243del (Beta) that occurs in the 74_Left primer, and the K417N (Beta) or K417T (Gamma) which occurs in the 76_Left primer^10,11^. The Delta/B.1.1.672 variant has also been associated with ARTIC v3 drop-out of primers 72R and 73L^12^.

Tools such as Nextclade^13^ can capture and report these sequence anomalies. Reports generated through Nextcalde includes detailed information on excess number of gaps, mixed bases, private mutations and frameshifts. However, there is no easy to use bioinformatics QC tool to further explore the mapped short reads with respect to positional codons, nucleotide and indel distribution in microbial genomes, or to support batch updates to consensus sequence. Moreover, such tools are often taxonomically limited and remain optimized for a select set of reference genome sets.

Here, we introduce SeqPanther, a Python application that provides the user with a suite of tools to further interrogate the circumstance under which these mutations occur and to modify the consensus as needed for non-segmented bacterial and viral genomes where reads are mapped to a reference. SeqPanther generates detailed reports of mutations identified within a genomic segment or positions of interest, including visualization of the genome coverage and depth. Our tool is particularly useful in the examination of multiple next-generation sequencing (NGS) short read samples. Additionally, we have integrated Seqpatcher^14^ that supports the merging of Sanger sequences into their respective NGS consensus.

### Implementation

We utilised Python (v3.9.9) as the base programming language to develop our pipeline. Pysam (a wrapper around htslib and the samtools package, 0.18.0) is the core module^15,16^ of the pipeline which allows exploration and manipulation of BAM files generated by sequence mapping tools to extract the distribution of reads. The tool also uses the click package (v8.0.3) to capture user inputs, Pandas (v1.3.5)^17^ to store and arrange data in data-frames and to generate tables, and Matplotlib (v3.6.2)^18^ for plotting read distributions and highlighting the changes in a given region. The outputs from the tool were visually validated using the Geneious Prime 2023.0.1 program (https://www.geneious.com/).

### Features

SeqPanther, is an easy to use and accurate command-line tool for generating statistics relating to amino acid altering information in the generated reads, and integrating relevant changes to used references. SeqPanther features a suite of tools that perform various functions including codoncounter, cc2ns, and nucsubs. Figure 1 shows inputs, outputs and dependencies between the tools.

**Figure 1:**
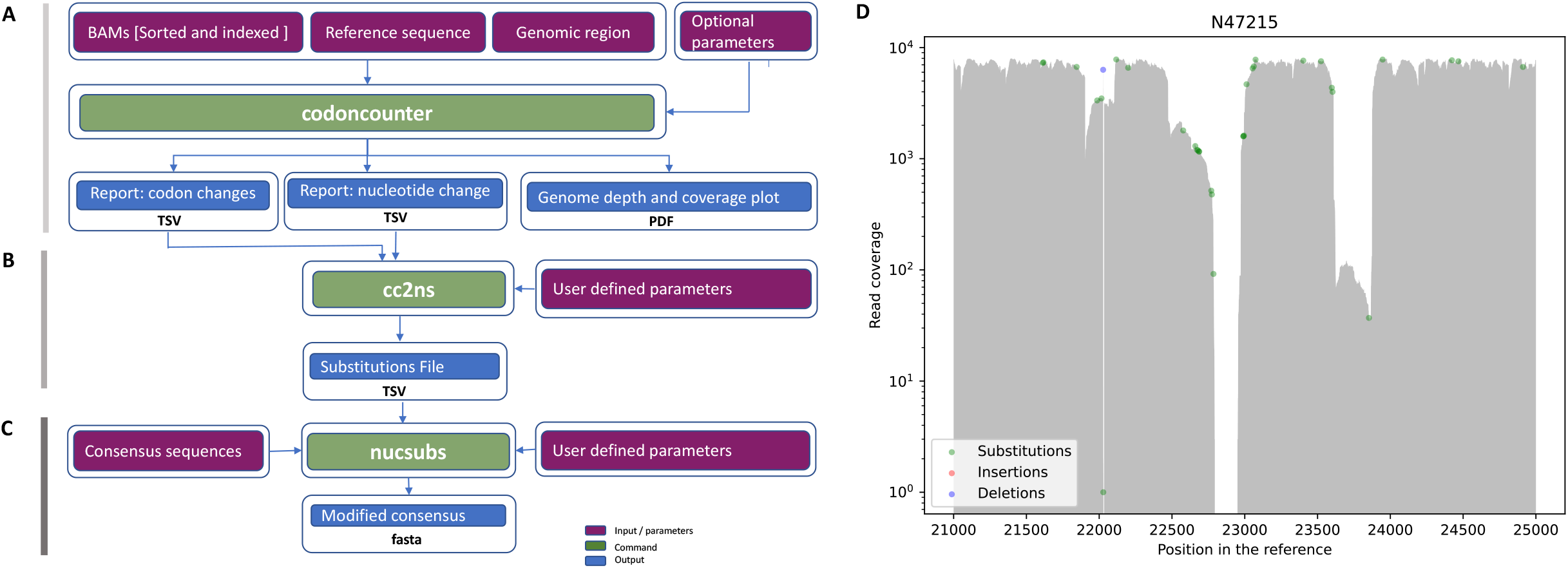
Visual illustration of the Seqpanther components. A) codoncounter which accepts a BAM file or directory containing BAM files (if running batch mode), a reference sequence and a genomic region or set of positions and generate two reports (codon and nucleotide) and a genome coverage and depth map. B) The cc2ns command accepts indel and substitution reports as inputs and generates a substitution file containing all the nucleotide substitutions and indels as well as their frequency. C) The nucsubs module that takes the file substitutions file, reviewed and edited by the user and modifies the consensus sequences. D) Depth and coverage plot generated by codoncounter. Colored circles represent different types of alterations across the length of the assembly, which are supported by more than 5% of total reads.

### Codoncounter

The *Codoncounter* module takes at least a bam file, reference fasta and gff genome annotation file as input to produces four output files: 1) a table summarizing codon variations impacting amino acid assignment in a protein sequence (Table S1); 2) a table listing the nucleotide substitutions (Table S2); 3) list of the nucleotide indels identified (Table S3); and 4) a PDF file with plots showing distribution of reads and types of alterations at different genomic coordinates that occur relative to the reference (Figure 1D). In case of unsorted and unindexed bam files, the command generates a temporary sorted and indexed bam. Changes in amino acid are displayed relative to the strand where the gene is present. While calculating the impact on amino acid, incase of substitution even, all three nucleotides must be aligned. Reads with indel even around a codon were not merged with substitutions and explored independently.

Additionally, for each output type generated by Codoncounter, the command provides additional statistics. These include the total number of reads, reads mapping to the reference, alternate substitutions and the percentage they comprise. Typically, the codon results are tied to a user specified region such as an open reading frame (ORF) or protein coding gene with coordinates defined in a general feature format (gff) file. The focus on a user specified genomic segment is based on the fact that some samples might have an extremely large mutation load and quality control will only be focused on just a small subset of mutations in a region of interest, for example, in the case of SARS-CoV-2 mutations in the Spike gene or in the Receptor Binding Domain (RBD). The user can also specify the set of nucleotide positions or a genomic range of interest, or both. The user provided reference sequence ID must match the ID in the bam, gff, and reference FASTA file. The codons report only contains non-synonymous substitutions or 3x nucleotide indels in the mapped reads (Supplementary Table 1). In comparison, all nucleotide changes are reported including the synonymous changes i.e. mutations not associated with an amino acid change (Supplementary Table 2). The report is thus also useful to investigate synonymous mutations which, although not change the encoded amino acid, play a crucial role in the estimation of the molecular clock ^19^ and designation of strains and lineages. Examples of lineage defining synonymous mutations include the T23341C and T24187A in the P3^20^ variant and C26801T in the Alpha/B.1.1.7 variant ^21^.

**Table 4:**
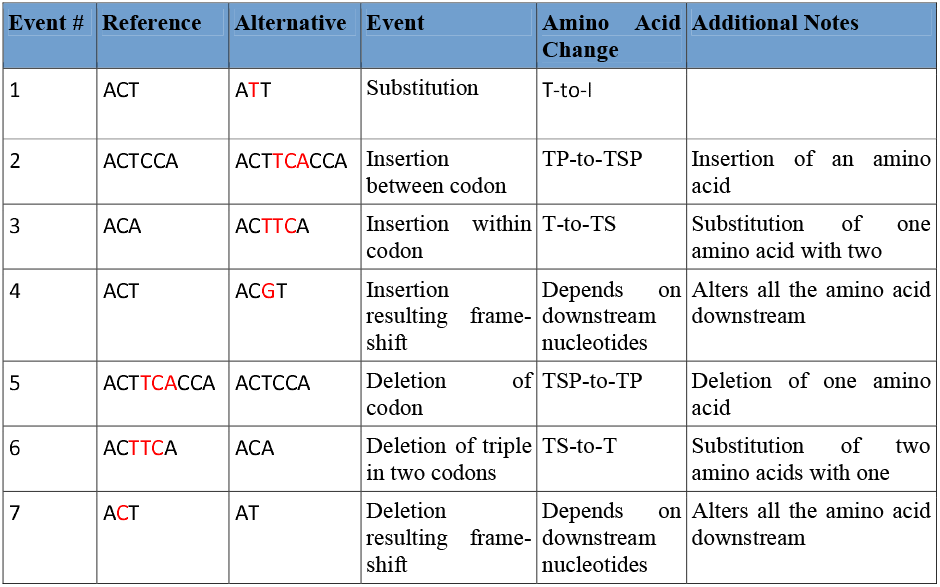
Types of changes resulting in amino acid alterations. In case of indels, only multiples of 3 consecutive nucleotides indels were considered for integration in consensus sequences.

The output from this module also provides insight into sub-consensus mutations that could potentially become fixed in the population. By default, the tool implements a threshold of 5% to report mutations, however this can be adjusted up or down to the desired detection threshold by the user. The tool can be run in batch mode or on a single BAM file. To execute the batch mode, the user simply provides a folder containing the BAM files to be analysed. The output tabular reports can be imported into other tools for further downstream analyses.

### Cc2ns

The *cc2ns* command takes the output from the *codoncounter* command (nucleotide substitution and indel reports) and generates a tabular file according to the user specified filters that contain the changes which could be integrated in the strain-specific consensus sequences. The output file contains one change per row. In case of multiple changes to a coordinate, the changes are provided in multiple rows. The user can edit the file by adding additional changes or by removing suggested alterations. In case of multiple changes at a position, the first instance is accepted. The reviewed file is passed to the *nucsubs* command to modify the consensus sequences. The file contains a substitution percentage (i.e. of the reads supporting the substitution). Although this value is optional, it can be used to specify a threshold for all mutations to be restored. This is particularly useful when mutations are not called due to higher thresholds in the variant calling pipeline and is recommended mostly for known, fixed mutations within a constellation.

### Nucsubs

The *nucsubs* command modifies consensus sequences by integrating the user-defined base substitutions, deletions and insertions by adding additional nucleotides at a given position relative to a reference sequence. As consensus sequences usually differ in length from the reference, to integrate the changes, coordinates are specified relative to the reference. To acquire relative alteration positions, consensus sequences are aligned to the reference using MAFFT aligner v7.508 ^22^ in auto mode. The MAFFT alignment is then imported into a pandas dataframe for coordinate manipulation and integration of changes. The modified sequences are saved as a FASTAfile. The functionality expedites the often manual process of modifying consensus sequences that often introduces new artifacts when modifying a large number of sequences. The module is sufficiently intelligent to only update the sequences of interest within the alignment or sequence file as required by the user.

These modules were tested on South African SARS-CoV-2 strains and the results validated manually with the Geneious Prime programme (version 2022.2.2). Manual editing of alignments is a common practice in molecular analyses to improve the quality of sequences prior to phylogenetic inference^23^.

### Seqpatcher

In addition to the new features, we have integrated Seqpatcher^14^, our bioinformatics tool to merge Sanger sequence fragments into NGS-based consensus sequences. Seqpatcher is particularly useful in the case of primer drop-outs resulting in significantly large gaps spanning an entire protein, open reading frame or gene. Seqpatcher accepts Sanger sequences both in FASTA (pre-processed) or chromatograph (raw .ab1) formats. Sanger sequencing to cover such gaps that are less than a thousand bases is effective in both cost and time relative to re-sequencing the whole genome.

## Conclusion

Genomic epidemiology and associated public health interventions entirely depend on the quality of pathogenic genomic data. False positive or negative sequencing outputs can have dire consequences on the downstream analyses and subsequent public health decisions. Quality control and assurance are therefore critical components of the sequence generation process. SeqPanther provides a unique suite of tools to explore short reads and modify consensus sequences. The functionalities provided by Seqpanther can significantly simplify the process of and improve the speed of sequence quality control in small to medium-sized sequencing laboratories, thus in turn reducing the turnaround time to provide quality genomic sequences. Although the tools have been primarily developed and tested using SARS-CoV-2 data, they can be applied to most non-segmented bacterial and viral genomes where reads are mapped to a reference.

## Future prospects

Many microorganisms have fragmented genes (Introns separated exons), overlapped genes or genes with frameshift events such as Influenza A with a segmented genome, however, at present the tool only supports single segment genomes without other stated events. We plan to extend it to support complex gene and genome structures. We wish to report indels whose length is not a multiple of 3 and yet they are present in the same read and summation leads to multiple of three. Furthermore, as the cost of sequencing continues to reduce and capacity to sequence increases, we expect to see a further influx in the number of microbial genomes. To this effect, we intend to implement performance enhancements to reduce the batch analyses running time given a fairly large number of genomes.

## Authorship contribution

Conceptualization: JES, TdO, AMK ^*τ*^; Methodology: HT, JES, AMK^*τ*^, TdO; Investigation: JES, AMK^*τ*^; Visualisation: JES, AMK^*τ*^; Funding acquisition: TdO; Project administration: TdO, JES, EW; Supervision: TdO; Writing – original draft: JES, AMK^*τ*^; Writing – review & editing: HT, JES, EW, TdO, SvW, SE, AMK, AMK^*τ*^

---

**Table S1:**
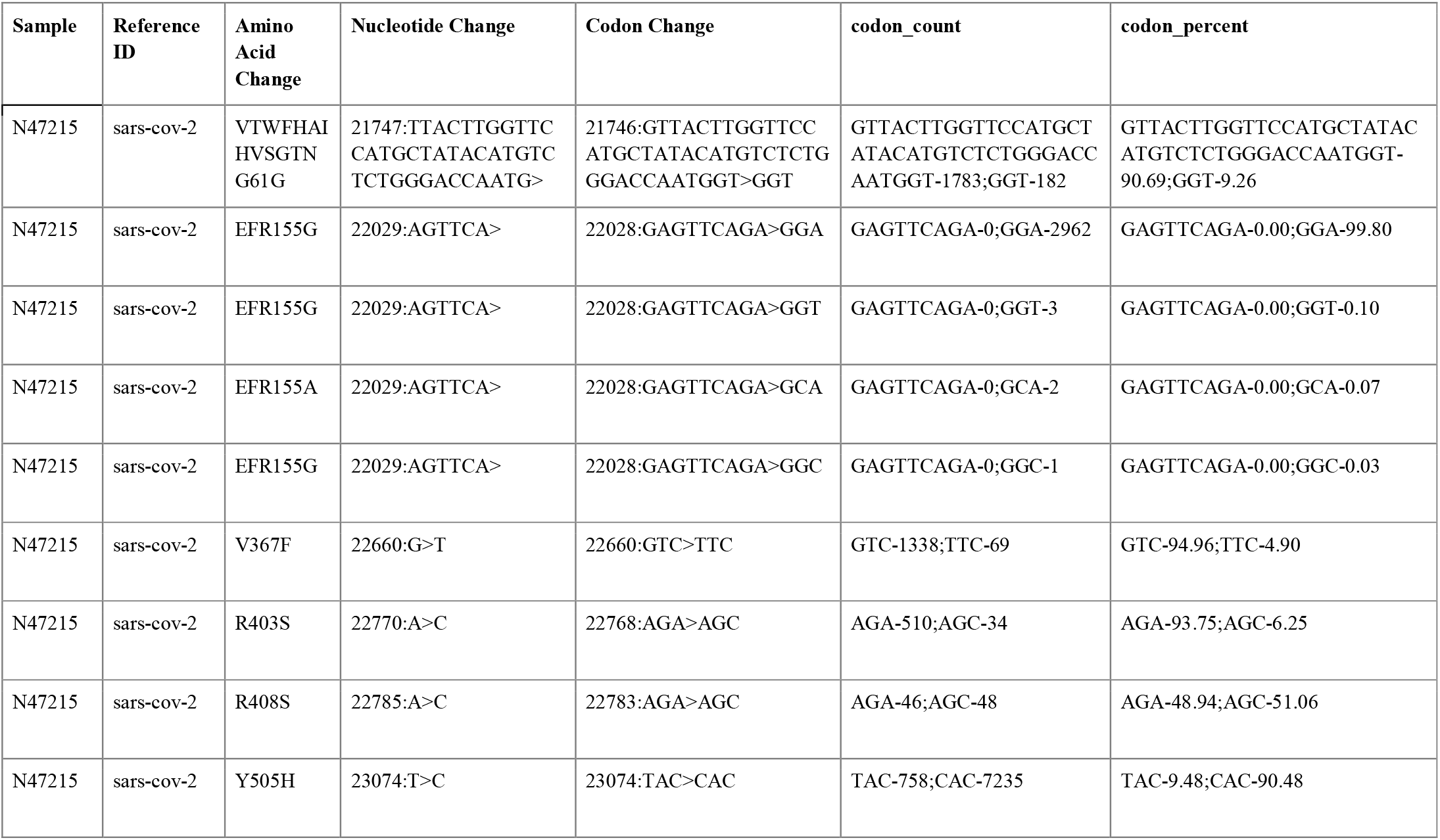
Codon changes affecting amino acid in sars-cov-2 spike-protein gene their distributions

**Table S2:** Nucleotide substitution in sars-cov-2 spike-protein gene their distributions

| Sample | Reference ID | pos | Reference Nucleotide | read_count | Nucleotide Frequency | Nucleotide Percent |
| --- | --- | --- | --- | --- | --- | --- |
| N47215 | sars-cov-2 | 21613 | C | 7271 | T:7261 | T:100 |
| N47215 | sars-cov-2 | 21617 | C | 7367 | G:7359 | G:100 |
| N47215 | sars-cov-2 | 21845 | C | 6698 | T:6688 | T:100 |
| N47215 | sars-cov-2 | 21986 | G | 3345 | A:3343 | A:100 |
| N47215 | sars-cov-2 | 22016 | G | 3488 | T:3486 | T:100 |
| N47215 | sars-cov-2 | 22117 | T | 7812 | C:7807 | C:100 |
| N47215 | sars-cov-2 | 22199 | T | 6581 | G:6574 | G:100 |
| N47215 | sars-cov-2 | 22577 | G | 1796 | A:1793 | A:100 |
| N47215 | sars-cov-2 | 22660 | G | 1301 | G:1241,T:60 | G:95,T:5 |
| N47215 | sars-cov-2 | 22673 | C | 1199 | T:1197 | T:100 |
| N47215 | sars-cov-2 | 22678 | T | 1191 | C:1189 | C:100 |
| N47215 | sars-cov-2 | 22685 | C | 1161 | T:1159 | T:100 |
| N47215 | sars-cov-2 | 22687 | A | 1161 | G:1161 | G:100 |
| N47215 | sars-cov-2 | 22770 | A | 516 | A:480,C:34 | A:93,C:7 |
| N47215 | sars-cov-2 | 22774 | G | 477 | A:477 | A:100 |
| N47215 | sars-cov-2 | 22785 | A | 92 | A:44,C:48 | A:48,C:52 |
| N47215 | sars-cov-2 | 22990 | A | 1592 | G:1592 | G:100 |
| N47215 | sars-cov-2 | 22991 | G | 1592 | A:1592 | A:100 |
| N47215 | sars-cov-2 | 22994 | C | 1616 | A:1616 | A:100 |
| N47215 | sars-cov-2 | 23012 | A | 4675 | C:4672 | C:100 |
| N47215 | sars-cov-2 | 23054 | A | 6498 | G:6495 | G:100 |
| N47215 | sars-cov-2 | 23062 | A | 6767 | T:6761 | T:100 |
| N47215 | sars-cov-2 | 23074 | T | 7791 | T:712,C:7075 | T:9,C:91 |
| N47215 | sars-cov-2 | 23402 | A | 7626 | G:7619 | G:100 |
| N47215 | sars-cov-2 | 23524 | C | 7536 | T:7524 | T:100 |
| N47215 | sars-cov-2 | 23598 | T | 4350 | G:4346 | G:100 |
| N47215 | sars-cov-2 | 23603 | C | 3995 | A:3988 | A:100 |
| N47215 | sars-cov-2 | 23853 | C | 37 | A:37 | A:100 |
| N47215 | sars-cov-2 | 23947 | G | 7800 | T:7778 | T:100 |
| N47215 | sars-cov-2 | 24423 | A | 7689 | T:7680 | T:100 |
| N47215 | sars-cov-2 | 24468 | T | 7529 | A:7508 | A:100 |
| N47215 | sars-cov-2 | 24911 | C | 6695 | T:6679 | T:100 |

**Table S3:**
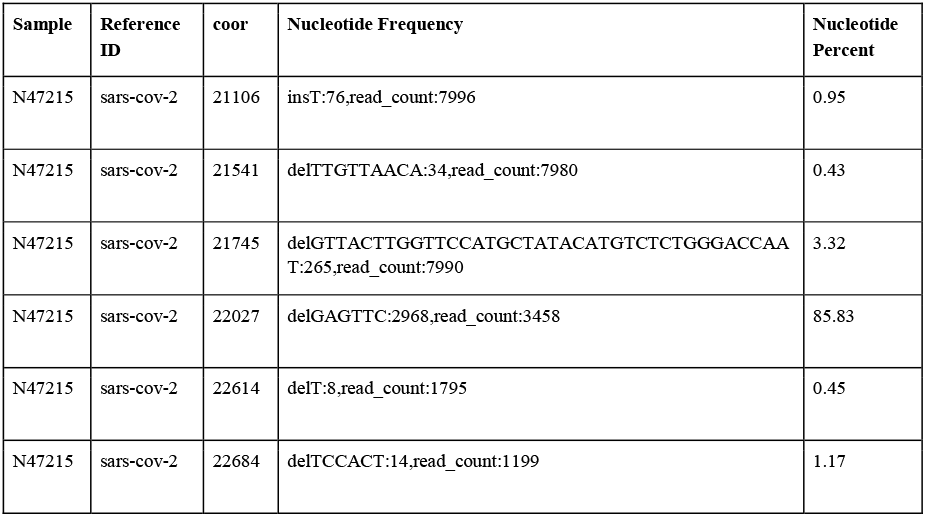
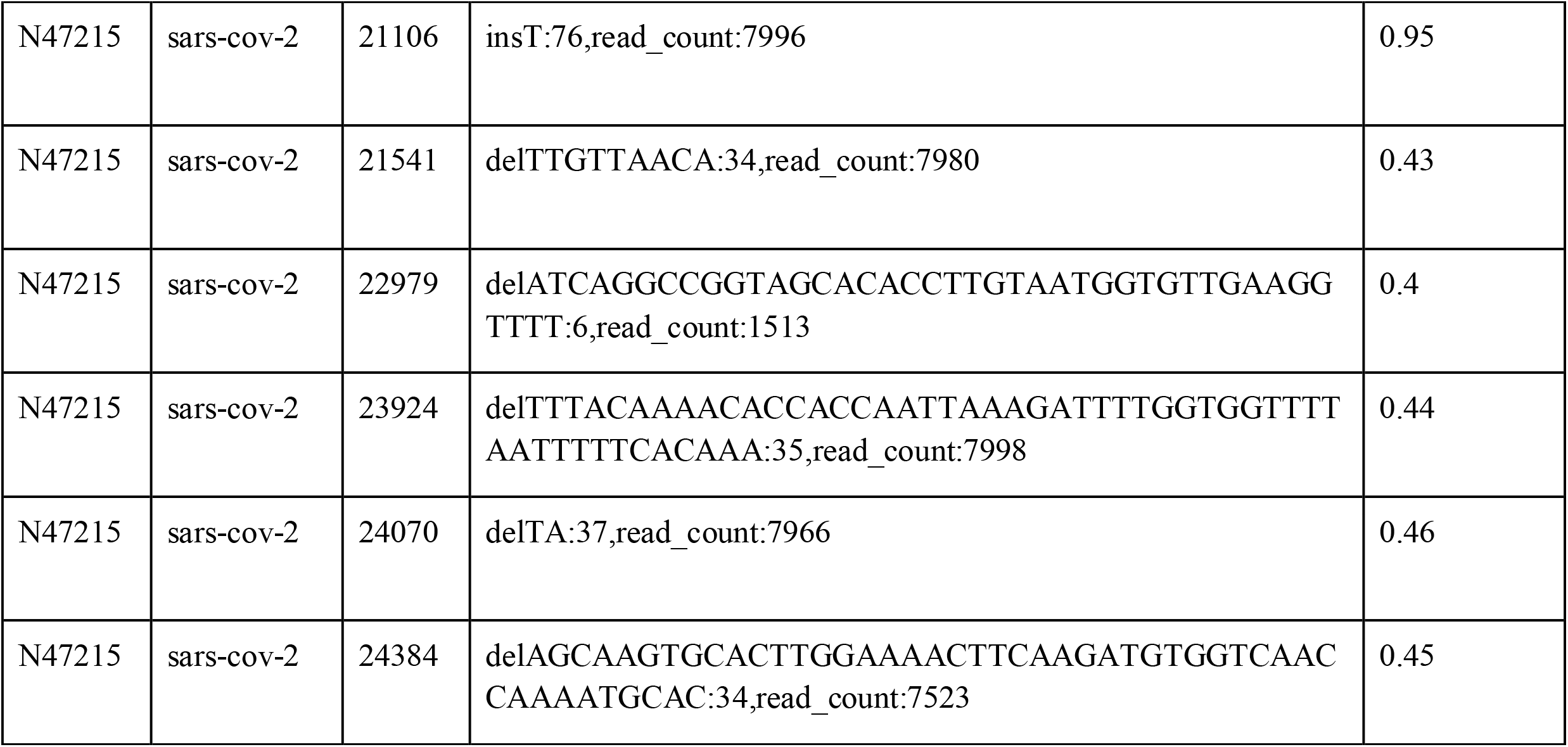
Selected nucleotide InDels in sars-cov-2 spike-protein gene their distributions

